# CTCF-mediated Genomic Effects of BART Region on Epstein-Barr Virus Chromatin 3D Structure in Gastric Carcinoma Cells

**DOI:** 10.1101/2020.12.03.409722

**Authors:** Kyoung-Dong Kim, Subin Cho, Taelyn Kim, Sora Huh, Lina Kim, Hideki Tanizawa, Jae-Ho Shin, Teru Kanda, Kyoung Jae Won, Paul M. Lieberman, Hyosun Cho, Hyojeung Kang

**Affiliations:** Department of System Biotechnolog, Chung-Ang University, Anseong, Korea; Vessel-Organ Interaction Research Center, VOICE (MRC), Cancer Research Institute, College of Pharmacy, Kyungpook National University, Daegu, Korea; Institute of Molecular Biology, University of Oregon, Eugene, Oregon, United States of America; Department of Applied Biosciences, Kyungpook National university, Daegue, Korea; Division of Microbiology, Faculty of Medicine, Tohoku Medical and Pharmaceutical University, Sendai, Japan; Biotech Research and Innovation Centre (BRIC), University of Copenhagen, Copenhagen, Denmark; Program in Gene Expression and Regulation, The Wistar Institute, Philadelphia, Pennsylvania, United States of America; Duksung Innovative Drug Center, College of Pharmacy, Duksung Women’s University, Seoul, Korea

## Abstract

EBV latent infection in gastric carcinoma (GC) cells is characterized by distinct viral gene expression programs. CCCTC-binding factor (CTCF) is a chromatin structural factor that has been involved in coordinated chromatin interactions between multiple loci of Epstein-Barr virus (EBV) genes. Here, we investigate the role of CTCF in regulating EBV gene expression and chromosome conformation in model of EBV-associated gastric carcinoma (EBVaGC). Chromatin immunoprecipitation followed by sequencing (ChIP-seq) against CTCF revealed 16 CTCF binding sites (BS) in EBV genome of EBVaGC, SNU719 cells. Among the CTCF BSs, one site named as BARTp (BamHI A right transcript promoter) CTCF BS is located at upstream of 11.8-kb BART region (EBV genome: 139724-151554) and was not yet defined its biological functions in EBV life cycle. EBV BART encodes a complex miRNA cluster of highly spliced transcripts that is implicated in EBV cancer pathogenesis. This present study investigated the functional role of the CTCF binding site at BARTp (BARTp CTCF BS) in regulating EBV gene transcription and EBV three-dimensional (3D) genome structure as DNA loop maker. Circular chromatin confirmation capture (4C)-seq and chromatin confirmation capture (3C)-semi-quantitative(sq)PCR assays using SNU719 cells revealed that BARTp CTCF BS interacts with CTCF BSs of LMP1/2, Cp/OriP, and Qp in EBV genome. We generated mutations in BARTp CTCF BS (S13) in bacmids with (BART^+^) or without (BART^−^) the 11.8-kb BART transcript unit (B(+/−)). ChIP-qPCR assay demonstrated that CTCF binding was ablated from BARTp in EBV B(+/−) S13^−^ genomes (mutant S13), elevated at several other sites such as *LMP1, OriP*, and *Cp* in EBV B(-) (BART^−^) S13^−^ genome, and decreased at the same sites in EBV B(+) S13^−^ genome. Infection assay showed that BARTp CTCF BS mutation reduced infectivity, while BART transcript deletion has no detectable effects. Gene expression tests showed that *EBNA1* was highly downregulated in B(+/−) S13^−^ EBVs related to B(+/−) S13^+^ EBVs (wild-type S13). *LMP1* and *BZLF1* were more downregulated in B(-) S13^−^ EBV than B(+) S13^−^ EBV. Taken together, these findings suggest that the CTCF binding and BART region contribute to EBV 3D genome structure via a cluster of DNA loops formed by BARTp CTCF BS (S13) and are important for coordinated viral gene expression and EBV infectivity.

## Introduction

Epstein-Barr virus (EBV) is a member of the human gamma herpesvirus family that establishes lifelong latent infection in the most population [1]. EBV latent infection is associated with lymphoma such as Burkitt’s lymphoma (BL) and Hodgkin’s lymphoma (HL), and epithelial neoplasm such as nasopharyngeal carcinoma (NPC) and gastric carcinoma (GC) [2, 3]. During the latency phase, EBV genome exists as multicopy episomes that express only a few of viral genes called latent genes [4]. The latency can be changed into the lytic phase depending on developmental stage, environmental signals and pharmacological manipulation [5, 6]. Approximately 10% GC has been diagnosed as EBV associated gastric carcinoma (EBVaGC), estimating more than 70,000 cases worldwide per year [7-9]. EBVaGC appears lymphoepithelioma-like carcinoma whose definition is an undifferentiated carcinoma with lymphocytic infiltrate, histologically similar to NPC [2, 7]. EBV of EBVaGC maintains the type I latency phase and express the narrowest group of EBV latent genes such as *EBNA1, EBER*, BARTs and sometimes *LMP2A*. These genes are implicated with the EBVaGC oncogenesis [10].

The EBV genome contains two miRNA cluster that encoded by BamHI fragment H rightward open reading frame 1 (BHRF1) and BamHI A right transcripts (BARTs) [11-14]. EBV BARTs are a complex miRNA cluster of highly spliced transcripts initially found in NPC EBV strain [15, 16]. Some lymphotropic BL EBV strain, like B95-8, have a deletion overlapping the 11.8-kb BART region (139724-151554), while EBV strains derived from GC, such as GC1 and YCCEL1, contain the full BART region [17, 18]. BART miRNAs are substantially expressed in EBV infected epithelial cells such as NPC and EBVaGC [19-21]. The BART miRNAs are highly implicated in EBV-mediated epithelial malignancies but sometimes dispensable in EBV-mediated lymphoma. Thus, their function in EBV life cycle is only partially elucidated [22].

The maintenance of chromatin structure is also largely dependent on cellular mechanisms that regulate several chromatin interactions exemplified an interaction between enhancer and promoter [23-26]. The CCCTC-binding factor, also referred to CTCF, is a transcription factor that contains DNA binding domain and 11 zinc fingers. CTCF is involved in other functions such as epigenetic insulator, gene boundary factor and DNA looping maker [27-29]. In particular, CTCF is highly associated with regulating long range chromatin interaction by chromatin loop organization [30]. Cohesin composed of SMC1, SMC3, and non-SMC components including RAD21, SA1, and SA3 are known to assist in CTCF-mediated stabilization of EBV genome structure [31-33]. Cohesion binds at multiple control regions of EBV genes and involves in maintain EBV genome structure to regulate EBV gene expression, along with CTCF [34-36].

Here, we have identified 16 CTCF binding sites (BS) in the EBV genome in EBVaGC using ChIP-seq analysis against CTCF. Among them, one site (BARTp CTCF BS) located in close proximity to the transcription start site of the 11.8-kb BART region that has not yet been defined for its biological function in EBVaGC. This CTCF binding site, referred to here as the BARTp CTCF BS (S13), exits in most EBV genomes regardless of the existence of 11.8-kb BART. Here, we test the hypothesis that the BARTp CTCF BS (S13) is important for 3D conformation of EBV genome, and that the BART transcripts affect this conformation. We further test whether BARTp CTCF BS (S13) regulates EBV gene expression. We find that BARTp CTCF BS (S13) contributes to both BART transcription regulation and 3D conformation, and is likely to contribute to regulation of EBV oncogenesis and life cycle.

## Materials and Methods

### Cell lines and Reagents

HEK293 cells and HEK293-EBV bacmid cell lines were cultured in Dulbecco’s Modified Eagle Medium (DMEM; Hyclone, Pittsburgh, PA, USA) supplemented with 10% Fetal Bovine Serum (FBS; Hyclone, Marlborough, MA, USA), antibiotics/antimycotics (Gibco, Waltham, MA, USA), and Glutamax (Gibco, Waltham, MA, USA) at 37°C, 5% CO_2_, 95% humidity in a CO_2_ incubator. HEK293 cells were transfected with EBV bacmid and them selected with hygromycin B (400 μg/ml) (Wako, Japan). Gastric carcinoma cell lines SNU719 (EBVaGC) was purchased from Korean Cell Line Bank (Seoul, Korea) and cultured in RPMI 1640 medium (Hyclone, Pittsburgh, PA, USA) supplemented with 10 % FBS, antibiotics/antimycotics and GlutaMAX at 37 °C with 5% CO_2_ and 95% humidity.

### Microscale thermophoresis (MST) assay

Wild type (Wt) S13 49-mer primer set spanning EBV 138,946~138,956 was designed whose forward primer was labeled with 5-carboxyfluorescein (5-FAM) (Fig. 2A). We in parallel designed mutant (Mt) S13 49-mer set as counterpartner for Wt S13 49-mer primer set. Mt S13 50-mer contains several point mutations in region spanning EBV 138,963~138,987 whose mutations were expected to disrupt CTCF binding to Wt S13 49-mer primer set (Fig. 2A). 10 μl of 100 μM forward Wt S13 49-mer primer was mixed with 10 μl of 100 μM reverse Wt S13 49-mer primer. The mixture was paired by placing it in a thermocycler which programmed to start at 95°C for 2 min and then gradually cool to 25°C over 45 min. In parallel, 10 μl of 100 μM forward Mt S13 49-mer primer was also mixed and pared with 10 μl of 100 μM reverse Wt S13 49-mer primer. Resultant paired both Wt and Mt (Wt/Mt) S13 49-mer primer sets were used as target DNA in MST assay. Using baculovirus expression system, his-tagged CTCF proteins were purified from Sf9 cells transfected with CTCF expression plasmid. Resultant CTCF proteins were used as ligand protein in MST assay. To make CTCF bind to Wt/Mt S13 49-mer primer sets, reaction mixture was prepared as followed: 5 ul 4X EMSA buffer (400 mM KCl, 80 mM HEPES, 0.8 mM EDTA, 80% glycerol, pH 8.0) freshly added 1 mM DTT, target DNA (125 nM), 1 ul 500 ng/ul sonicated salmon sperm DNA, 2.5 ul ligand protein (CTCF, 3.12 μM, 6.25 μM), and 8.5 ul sterile water. Afterward, each mixture was incubated 30 min on ice and subjected to Nanotemper Monolith NT.115 (Munich, Germany) as recommended by manufacturer.

**Fig 2.**
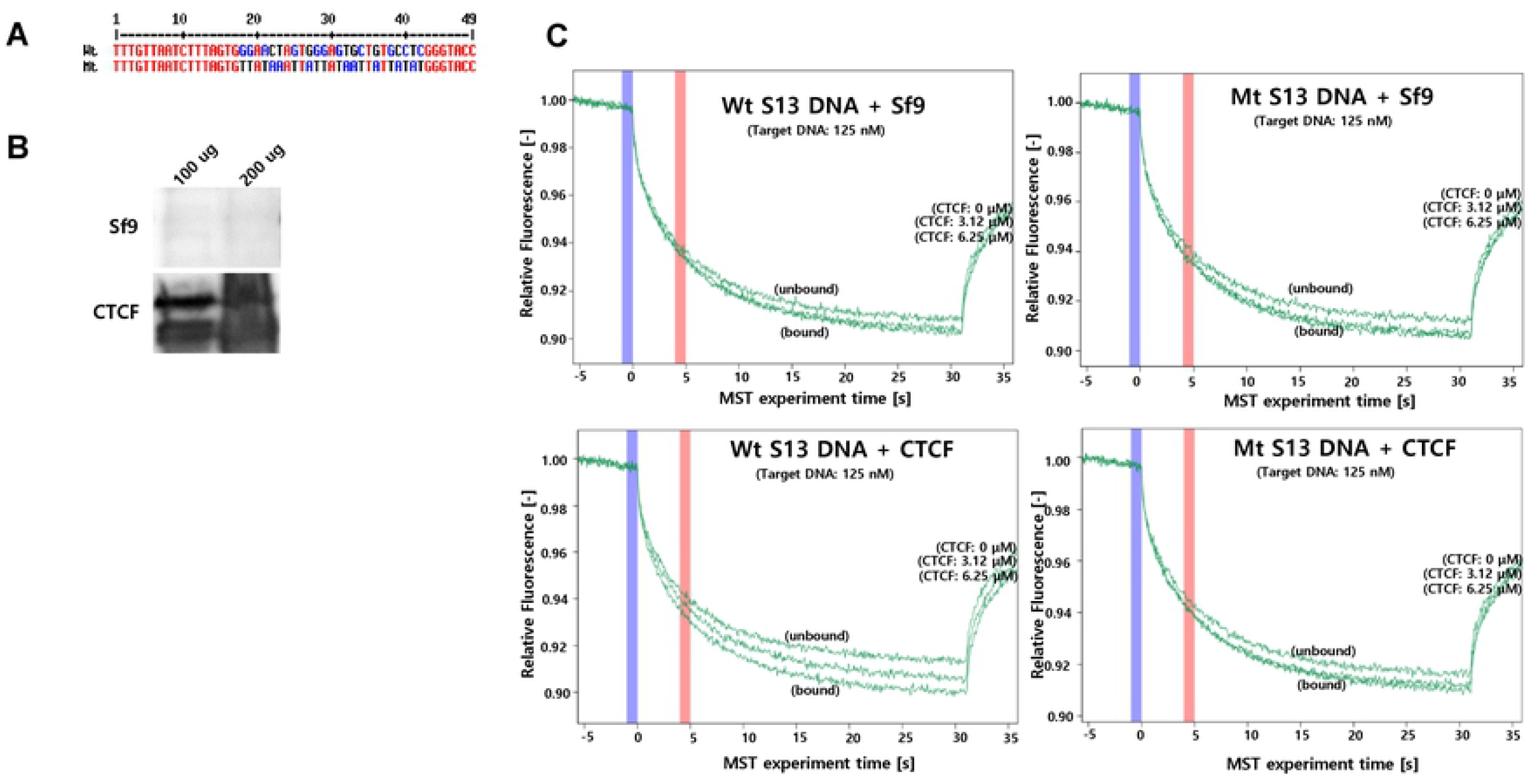
Detection of CTCF-DNA interaction using the Monolith NT.115. **A)** S13 5’ FAM labeled 49-mer primer whose sequence was mutated (Mt S13 primer sets) or not (Wt S13 primer sets). **B)** Confirmation of CTCF quality by Western blot assay. CTCF was purified using Sf9 baculovirus protein expression system. Upper panel: Sf9 total protein, lower panel: purified CTCF protein. **C)** CTCF-DNA interaction tested by Microscale thermophoresis (MST) assay. Wt/Mt S13 5’ FAM labeled 49-mer primer sets were paired to convert double DNA fragments. MST assays for CTCF binding to Wt/Mt S13 DNA fragments were conducted with 125 nM DNA fragments and 3.12 μM or 6.25 μM CTCF.

### PCR

To confirm BART region in EBV genome, PCR was performed at BART region using each EBV BART^+^ (B(+)) and EBV BART^−^ (B(-)) S13^−^ bacmids. One directional primers were used to avoid the amplication for self-ligation products; primers for PCR products of BART region from EBV B(+) Wt bacmid were used the forward primer. Each 25μl reaction contained 5 μl of EBV B(+) and B(-) Wt bacmid templates, 5 μl of 5× reaction mix (NanoHelix, Korea), 5 μl of 5× TuneUp solution (NanoHelix, Korea), 1 μl of Taq-plus polymerase (NanoHelix, Korea), and 2.5 μl of 10 μM forward/reverse primer. The following cycle conditions were used: 95°C for 3 min; 30 cycles of 95°C for 30 s, 55°C for 30 s, and 72°C for 30 s; followed by 72°C for 10 min. The reactions were performed using a TaKaRa PCR Thermal Cycler (Otsu, Japan) and then run on a 1.5% agarose/TBE gel.

### Real-time quantitative PCR (qPCR)

Quantification of precipitated DNA was determined using real-time quantitative PCR (qPCR) with SYBER Green in FastStart Essential DNA Green Master (Roche, Basel, Switzerland). Each resultant DNA was diluted in nuclease-free water and was analyzed in triplicated for EBV-associated genes and CTCF binding sites. The PCR reaction mixture of 20 μl contained 5 μl of template DNA, 0.5 μM of each primer and 10 μl of Master Syber Green 1 mix (Roche, Basel, Switzerland). The Primer sequences listed in Table 1. The following cycles thermal conditions were used: 95°C for 10 min; 45 cycles of 95°C for 10 sec, 55°C for 10 sec, and 72°C for 10 sec; 95°C for 5 sec; followed by 65°C for 1 min. In case of necessity, semi-quantitative PCR (sqPCR) was also conducted as previously described [37]. PCR products were amplified in a 25 μL reaction solution containing 5 μL of 5× reaction mix, 5 μL of 5× TuneUp solution, 1 μL of Taq-plus polymerase, and 2.5 μL of 10 pmol forward/reverse primer. The following cycle conditions were used: 95°C for 3 min; 30 cycles of 95 °C for 10 s, Tm (specific to primer sets) for 30 s, and 72 °C for 30 s; followed by 72 °C for 10 min. The reactions were performed using a TaKaRa PCR Thermal Cycler (TaKaRa, Kyoto, Japan) and then run on a 1.2%agarose/TBE gel.

### Construction of recombinant EBV bacmid

Site-directed mutations were introduced at BARTp CTCF BS (S13) in EBV B(+) or B(-) (B(+/−)) bacmids to make EBV B(+/−) S13^−^ bacmids that contains mutation in BARTp CTCF BS. EBV B(+) are isogenic and made from EBV B(-)bacmid [38]. EBV B(+/−) S13^−^bacmids were generated using two-step red-mediated recombination method. Primers for PCR products of the kanamycin resistant gene (Kan^r^) from pEPKanS3 plasmid used the forward primer 5’-GCA TCT TTC TAA CCA GTA GGG GCC TCC ACC TAG GTG CTT TGT TAA TCT TTA GTG TAT ATA TAT ATA TAT ATA TAT ATA TGG GTA CCC CTA TCC TAC AAC CAA TTA ACC AAT TCT GAT TAG-3’and the reverse primer 5’-ACA GGG ATT ATC AAG ACA AGG AGC TCC GGT AGG ACC TAT AGG ATA GGG GTA CCC ATA TAT ATA TAT ATA TAT ATA TAT ACA CTA AAG ATT AAC AAA GGA TGA CGA CGA TAA GTA GGG ATA-3’. Resultant PCR products were composed Kan^r^ gene with I-SceI site, flanked by 60 bp downstream and upstream of EBV BARTp CTCF BS sequence surrounding the designed CTCF mutation sequence. These PCR products were electroporated into GS1783 competent cells containing EBV B(+/−) S13^+^ bacmids for 1^st^ round homologous recombination. EBV B(+/−) S13^−^ bacmid with Kan^r^ gene was recovered by positive selection, characterized by restriction enzyme digestion and transformed into GS1783 I-SceI-inducible competent cells. Kan^r^ gene was removed from EBV bacmid-Kan^r^ by 2^nd^ round homologous recombination and negative selection to make complete EBV B(+/−) S13^−^ bacmids. Final CTCF mutation was confirmed by restriction enzyme digestion and DNA sequencing of the homologous recombination site in EBV B(+/−) S13^−^ bacmids.

### Transfection

EBV B(+/−) S13^+/−^ (B(+)S13^+^, B(+)S13^−^, B(-)S13^+^, B(-)S13^−^) bacmids were transfected into HEK293 cells using Neon transfection system (Invitrogen, Carlsbad, CA, USA). 5 x 10^4^ cells were resuspended in 100 μl of serum-free media containing 5ug of each mutant recombinant bacmid. The electroporation was conducted with Neon electroporator (Invitrogen) set at 1350 V, 30 ms and 1 pulse. Cells were placed in media supplemented with 10% FBS for 48 h post-transfection.

### Western blotting assay

Western blotting was performed in HEK293-EBV B(+/−) S13^+/−^ (B(+)S13^+^, B(+)S13^−^, B(-)S13^+^, B(-)S13^−^) cells. Cells (5 × 10^6^) were then lysed using 100 μl of RIPA lysis buffer (Promega, WI) supplemented with 1 μl of proteinase inhibitor and 10 μl of phenylmethylsulfonylfluoride. The cell lysates were further fractionated using the Bioruptor sonicator (5 min, 30 sec on/off pulses). Cell lysates were loaded onto 8% sodium dodecyl sulfate polyacrylamide electrophoresis gel and subjected to Western blot analysis by using antibodies against EBV proteins (1:1000 dilution). Following antibodies were used: anti-EBV EBNA1 (Santa Cruz Biotechnology (SCB), Santa Cruz, CA, USA), anti-EBV BZLF1 (SCB), EA-D (SCB), anti-LMP1 (SCB) and anti-GAPDH (Cell Signaling Technology, Danvers, MA, USA), Horseradish peroxidaseconjugated sheep anti-mouse IgG (Genetex, Irvine, CA, USA), horseradish peroxidase conjugated donkey anti-rabbit IgG (Genetex), and horseradish peroxidase-conjugated goat anti-rat IgG (Bethyl Laboratories, Montgomery, TX, USA) were used as secondary antibodies.

### Immunofluorescence Assay

HEK293-EBV B(+/−) S13^+/−^ (B(+)S13^+^, B(+)S13^−^, B(-)S13^+^, B(-)S13^−^) cells were grown on coverslips in 24-well plates (2 × 10^5^/well). Next day, cells were fixed with 4 % paraformaldehyde for 30 min and were permeabilized with 0.25 % Triton X-100 in PBS for 90 min. Treated cells were blocked using 1 % BSA in PBS containing 0.1 % Tween 20 for 60 min. Sample were stained with EBNA1 antibody (1 : 40). After overnight incubation at 4 °C, coverslips were washed 3X in PBS and treated with Alexa-594 (Thermo Fisher Scientific, Waltham, MA) for 2 h at 4 °C. Alexa-594 was used to detect EBNA1. After washing 3X in PBT (PBS containing 0.5 % Triton X-100), coverslips were mounted with DAPI (SouthernBiotech, Birmingham, AL). Samples were analyzed using immunofluorescence confocal microscopy.

### CTCF ChIP-seq analysis

ChIP-seq against CTCF was performed with 5 x 10^7^ SNU719 cells per sample and crosslinked using formaldehyde. Crosslinked SNU719 cell lysates was sonicated to achieve a DNA fragment length of ~ 100 – 500 bp. Immunoprecipitation was carried out was carried out with 10 μg of either rabbit anti-CTCF (Millipore, Burlington, MA, USA) or control rabbit IgG (SCB), incubated overnight with antibody-coated Dynabeads protein A/G (Invitrogen). Incubated beads were washed with ChIP-seq wash buffer (50 mM HEPES, pH 7.5, 500 mM LiCl, 1 mM EDTA, 1% NP-40, 0.7% Na-Deoxycholate, 1x protease inhibitors) for 5 times, then washed once with 50 mM NaCl in TE buffer. Immunoprecipitated DNA was eluted with ChIP-seq elution buffer (50 mM Tris-HCl, pH 8, 10 mM EDTA, 1% SDS), reverse-crosslinked at 65°C, treated with RNase A (0.2 mg/ml) and proteinase K (0.2 mg/ml), purified with phenol and chloroform, then subjected to qPCR validation. Validated ChIP samples were isolated by agarose gel purification, ligated to primers, and then subject to Illumina-based sequencing using manufacturer’s protocol (Illumina, San Diego, CA, USA). ChIP-seq reads were mapped to the EBV wide-type reference genome (NC 007605) using Bowtie. For peak calling findPeaks command in Homer software was applied [39].

### ChIP assay

CTCF or Cohesin ChIP assays were performed with 3 x 10^6^ SNU719 cells or HEK293-EBV B(+/−) S13^+/−^ (B(+)S13^+^, B(+)S13^−^, B(-)S13^+^, B(-)S13^−^) cells per sample according to the cross-linking chromatin immunoprecipitation (X-ChIP) protocol provided by Abcam (Cambridge, UK) with a slight modification. The Bioruptor (BMS, Korea) was used to sonicate genomic DNA according to the manufacturer’s protocol. Sonicated cell lysates were subjected to immunoprecipitation with antibodies to CTCF (Millipore), SMC1 (Bethyl Laboratories, Montgomery, TX, USA), SMC3 (Bethyl Laboratories) and normal rabbit IgG (SCB). The precipitates were incubated with ChIP elution buffer (1% SDS, 100 mMNaHCO3); then the samples were reverse-crosslinked at 65°C overnight and purified on Promega columns. The purified DNA was analyzed using RT-qPCR. ChIP values were calculated as fold increases over the isotype specific IgG values for each primer sets.

### 3C and 4C-seq assays

Chromatin confirmation capture (3C) and circular chromatin confirmation capture (4C)-seq assays were performed as previously reported [40]. Briefly, 1 x 10^7^ SNU719 cells were fixed in 1% paraformaldehyde for 10 min at 37 °C. Nuclei were permeabilized by incubation with 0.5% SDS at 62 °C for 10 min. A half of DNA was digested with 100 units of *XhoI* (New England Biolabs (NEB), Ipswic, MA, USA) and ligated in the nucleus followed by in situ 3C protocol [41]. Other half of DNA was digested with 100 units of MboI (NEB) and ligated in the nucleus followed by in situ Hi-C protocol [42]. After reversal of crosslinks, 3C DNA was prepared and examined the association between two genomic loci by using 3C primers. For 4C-seq, DNA was digested with 100 units of MboI (NEB) and ligated in the nucleus followed by in situ Hi-C protocol [42]. 3C DNA was further digested with 100 units of Csp6I (NEB) and re-ligated. For each primer viewpoint, a total 10 to 100 ng DNA was amplified by PCR. All samples were sequenced with NovaSeq 6000 100 bp paired read. 4C-seq experiments from all viewpoints were carried out in biological replicates for SNU719 cells.

### 4C-seq data analysis

The 100-bp sequence paired-end reads were trimmed and grouped by cutadapt (version 2.10) based on the sequence from 4C bait location. Reads were aligned to EBV genome (NC_007605) using Bowtie2 (version 2.2.3) with iterative alignment strategy. Reads with low mapping quality (MapQ < 10) and reads that mapped to human repeat sequences were removed. Total aligned reads for each *i*-th position of non-overlapping 10-kb window (*N_i_*) were calculated. Then, converted to the *P*-values using the Poisson formula:

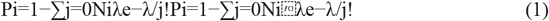

where *λ* is equal to the average of reads for each 10-kb window (except EBV aligned reads). The significant peaks were defined using subcommand bdgpeakcall of MACS2 software (version 2.1.1) with parameters: at least *P*-value < 10^-5^ (option −c 5), minimum length of 20-kb (option −l 20000) and maximum gap of 10-kb (option −g 10000). Total significant peak number for each 10-kb of bait positions (2 BART, FR, DS/Cp, Qp, LMP) were counted.

### EBV infection study

6.3 x 10^5^ HEK293-EBV B(+/−) S13^+/−^ (B(+)S13^+^, B(+)S13^−^, B(-)S13^+^, B(-)S13^−^) cells were seeded on 6-cm culture dish. On next day, HEK293-EBV B(+/−) S13^+/−^ cells were transfected with pCDNA3-BZLF1 and pCDNA3-BALF4 using TurboFect Transfection Reagent (Thermo Scientific, Waltham, MA, USA). Old medium was changed with fresh medium in one day post transfection. Medium was harvested in three days post transfection. Harvested medium was then clarified by twice centrifugation at 3000 rpm and once filtering with 0.45 μL filter (Sartorius stedium biotech, Gottingen, Germany). Next, 3 x 10^5^ HEK293 cells were plated on a well of 6-well culture plate. In next day, old medium was removed from the HEK293 cells. 40 μm cell strainer (FALCON, Corning, NY, USA) was loading on the HEK293 cells and refilled with the clarified old medium harvested from HEK293-EBV B(+/−) S13^+/−^ cells. EBV infection was occurred from the clarified medium to HEK293 cells through cell strainer for 24 h.

### Statistical Analysis

Statistical tests were performed using unpaired t-test and ANOVA. P-values (one-tailed) <0.05 (95% confidence) were considered statistically significant.

## Results

### Distribution of CTCF binding site (BS) in BART^+^ EBV genome

ChIP-seq was performed to identify CTCF BS in the EBV genome of EBVaGC SNU719 cells. The sequencing reads were mapped to the EBV reference genome (NC_007605) and were visualized using the Homer tools [39]. Approximately 5.7 × 10^4^ CTCF ChIP reads and 2.7 × 10^6^ IgG reads were aligned to EBV reference genome sequence and displayed reads on EBV genome using UCSC genome browser (Fig 1A). Enrichment of CTCF ChIP products relative to IgG ChIP products was calculated and visualized as the lower bound of 95% confidence. 16 high-confidence peaks were identified based on a read depth. CTCF binding sites were enriched at *BNRF1* (S1), *BCRF1* locus (S2), *BPLF1* locus (S5), *BMRP1* locus (S8), *BRLF1* locus (S11), *BART* (*RPMS1*) locus (S13), and *LMP1/2* locus (S16). These peaks were located in control regions of EBV genes such as upstream or downstream of their transcriptional start sites. One peak, S13 is located close to the BART promoter region and within the intro of *RPMS1* gene. Given to its locations, we speculate that this CTCF binding site (S13) functions as a transcriptional regulator and/or DNA loop maker, which mediate transcriptional regulator via chromatin interactions in EBV genome. To confirm CTCF BSs identified from ChIP-seq analysis in SNU719 cells, we performed ChIP-qPCR assays with CTCF and cohesion antibodies in these same cells. Since cohesins often colocalize with CTCF sites [32], we also assayed cohesion binding at these sites (Fig. 1B and 1C). CTCF and SMC1 strongly bound to most (four sites among tested five sites) CTCF BSs identified from ChIP-seq, while SMC3 was weakly bound to the CTCF BSs tested. In particular, CTCF and cohesins were more strongly recruited to S13 compared with other tested regions except S16. Taken together, all these results speculated that CTCF bound to S13 is likely to play dominant roles in chromatin interactions for EBV gene regulation in SNU719 cells.

**Fig 1.**
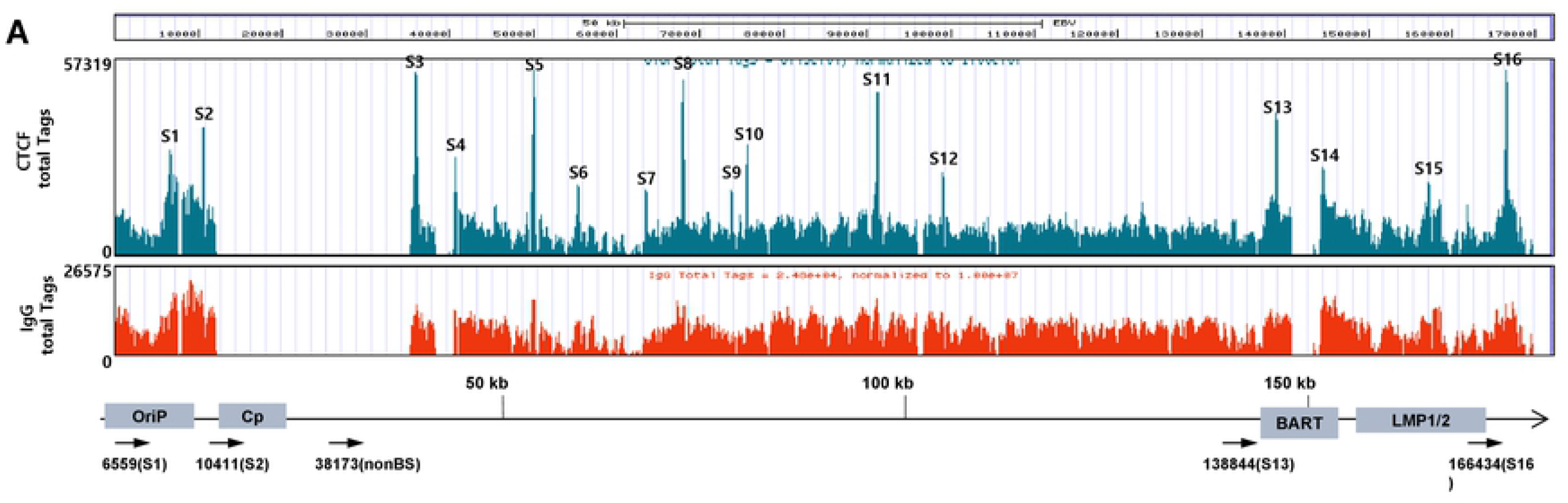
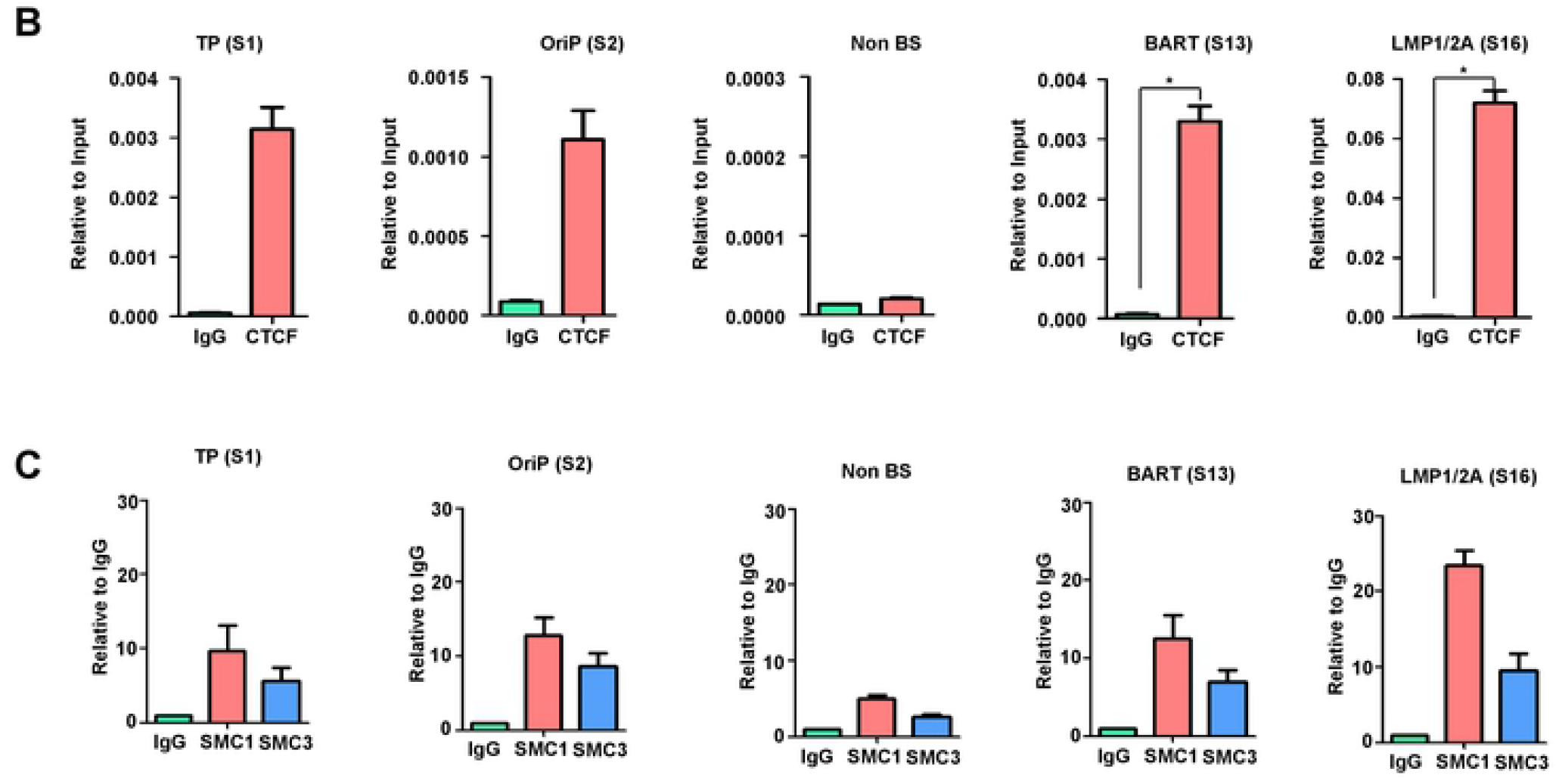
Identification of CTCF binding site on EBV genome in SUN719 cell line. **A)** CTCF enrichment in the EBV genome of SNU719 cells was identified by CTCF ChIP-seq assay. ChIP-seq data in SNU719 cells were aligned to EBV reference genome (NC_007605) for CTCF enrichment. Peak calling identifies 16 significant peaks in ChIP-seq signals. Approximate positions in EBV genome were indicated in the schematic with CTCF binding sites and primer sets used for ChIP RT-qPCR. **B)** Resultant CTCF enrichments identified by ChIP-seq assay were confirmed through the CTCF ChIP-qPCR assay. The ChIP-qPCR was assayed with SNU719 cells using antibody for CTCF. EBV 38,173 primer set was used as negative control. **C)** Like CTCF ChIP-qPCR assay, cohesin subunits such as SMC1 and SMC3 were also tested their enrichment at CTCF BSs by the cohesin ChIP-qPCR assays.

### The CTCF binding site in BART promoter region

Functional roles of CTCF BSs in EBV genome have been identified in EBV-associated lymphoid cells, such as Mutu I, Mutu-LCL, and Raji. CTCF BSs such as S1, S2, S5, and S16 in Fig.1A were characterized in previous studies and implicated in forming looping structure [41, 43, 44]. However, S13 (BARTp CTCF BS) was not previously characterized. To understand the role of CTCF binding at S13, we first validated that S13 bound directly to a CTCF protein from Sf9 cells using baculovirus CTCF expression system (Fig. 2B). To validate the specific sequence recognition site for CTCF, we used site-directed mutation to introduce mutations predicted to disrupt CTCF binding to S13 (Fig. 2A). We conducted microscale thermophoresis (MST) assay to confirm CTCF binding to wild-type (Wt) S13 and mutated (Mt) S13 DNA fragments (Fig. 2C). We observed strong binding of CTCF to Wt S13 DNA fragments that were significantly compromised for binding to Mt S13 DNA fragments (Fig. 2C lower panel). As control, total proteins isolated from Sf9 cells showed no strong selectivity in binding to Wt S13 or Mt S13 DNA fragments (Fig. 2C upper panel). These data indicated that Wt S13 is a good target site for CTCF binding and suitable for BARTp CTCF BS.

### Chromatin interaction between CTCF BS in BART^+^ EBV genome

To examine if BARTp CTCF BS (S13) is required for the formation of DNA looping structure, 4C-seq was conducted at around BARTp CTCF BS in SNU719 cells. Nucleus isolated from SNU719 cells were permeabilized, DNA-digested, and ligated to make 4C-seq products. Amplicons of 4C-seq products from view point primers were sequenced followed by aligning resultant sequencing reads to EBV genome (Fig. 3A). This 4C-seq data showed that the specific CTCF BS at S13 was involved in several chromatin interactions with multiple loci (Fig. 3B and 3C). Since CTCF binding site S14 is also located in the BART locus, we included S14 interactions along with S13. We found that BARTp CTCF BS (S13-S14, *RPMS1*) interacted strongly with several regions of the EBV genome, including regions at the EBV genome coordinates 5-kb region (S1, *BNRF1*=TP), 45-kb region (S4, *BPLF1*), 105-kb region (S12, *BKRF4*), and 155-kb regions (S15, *BALF4*) on the EBV genome. In particular, BARTp CTCF BS showed relatively strong interactions with both 45-kb region (S4) and 105-kb region (S12) except neighboring 155-kb region (S15). We also assayed interactions with other EBV regulatory elements using 4C-seq. OriP CTCF BS (S2) showed the strongest interactions its neighboring region at 5-kb (S1) and at 45-kb (S4). Qp CTCF BS (S5) interacted strongly with 145-kb region (S14), and LMP1/2 CTCF BS (S16) showed the strongest interaction with 55-kb region (S6). Duplicate experiments produced highly reproducible findings (Fig. 3C). These mutual chromatin interactions were depicted as a simple diagram (Fig. 3D). These results indicated that CTCF sites mediate DNA interactions throughout the EBV genome, and that BARTp CTCF BS intensively forms a cluster of several DNA loops with other important loci in the EBV genome of SNU719 cells.

**Fig 3.**
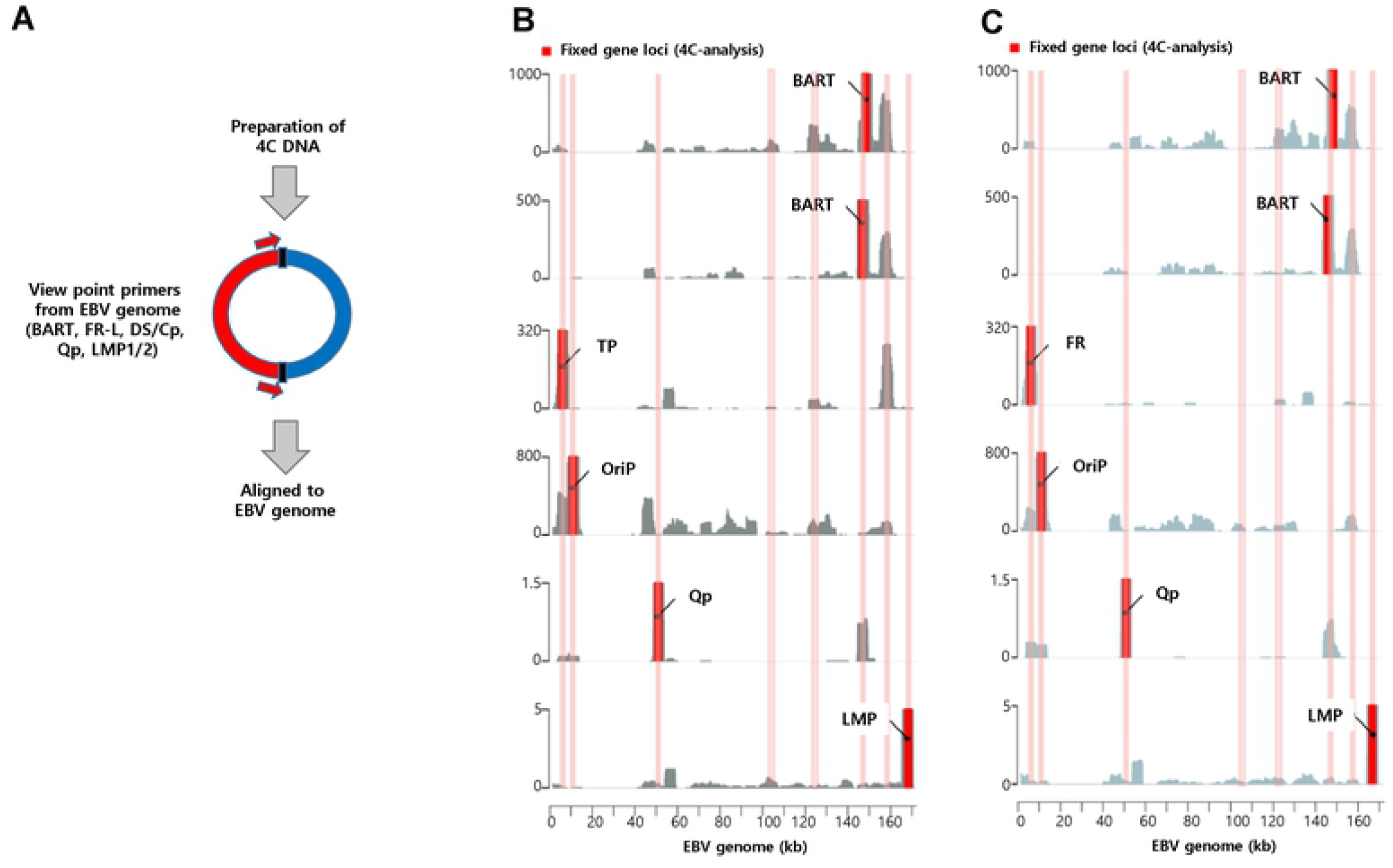
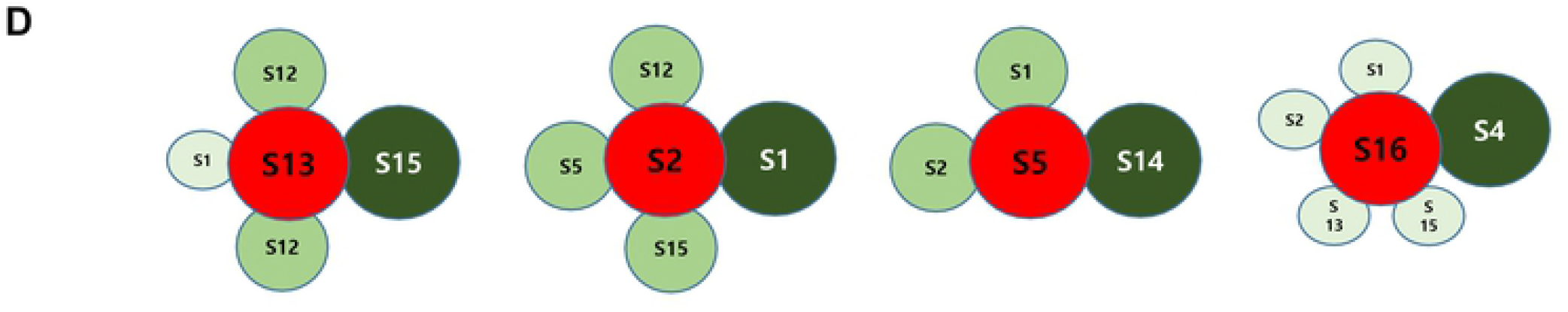
4C-plots on EBV genome. **A)** The brief 4C experimental procedure and bait positions on EBV genome. Five genomic loci (2 BART, FR, DS/Cp, Qp, LMP) were used as baits. **B)** and **C)** The entire EBV genome was divided into 10-kb windows with each 10 bp sliding, and read numbers indicating associations between the bait regions and 10-kb sections were plotted (top). The experiments are duplicated. **D)** Simple summary of multiple chromatin interactions mediated by CTCF BSs, given to 4C-seq assay data.

### Confirmation of chromatin interactions mediated BARTp CTCF BS

3C assay was conducted to consolidate diverse chromatin interactions identified by 4C-seq. Nucleuses isolated from SNU719 cells were subjected to paraformaldehyde-fixation, *XhoI*-digestion, T4 DNA ligase-ligation, and (nested) PCR. BARTp CTCF BS was included in an EBV DNA fragment cut by *XhoI* at 135,936 bp (135K, S13) and 147,676 bp (147K, S14) in EBV genome. Both ends of the 135K-147K DNA fragment were used as view point primers in PCR with 3C products (Fig. 4A). A view point primer designed from 147K (S14) in forward direction was first tested for chromatin interactions with other important loci in EBV genome (Fig. 4B). 147K (S14) locus could clearly interact with 3K (S1, *BNRF1*), 49K (S5, *BFRF3*), and 167K (S16, *LMP1/2*) loci. A second view point primer designed from 135K (S13) in reverse direction was found to interact with 65K (S8, BORL2) and 167K (S16, LMP1/2) loci. All chromatin interactions were summarized in table (Fig. 4D) and simply depicted as a simple diagram (Fig. 4E). Taken together, all these 3C data indicated that BARTp CTCF BS is involved in forming a cluster composed of at least three DNA loops with OriP CTCF BS (S2), Qp CTCF BS (S5), and LMP1/2 CTCF BS (S16), respectively.

**Fig 4.**
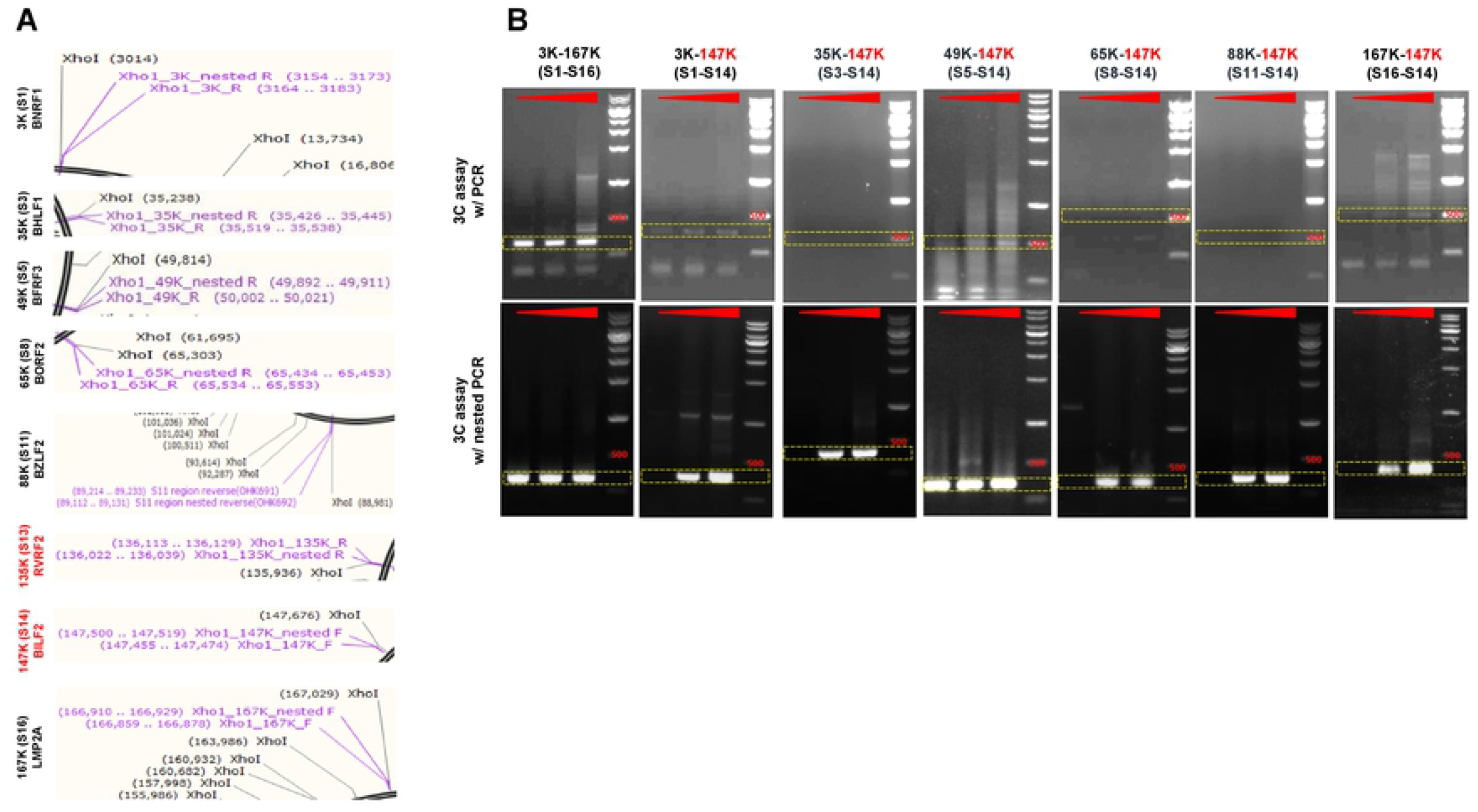
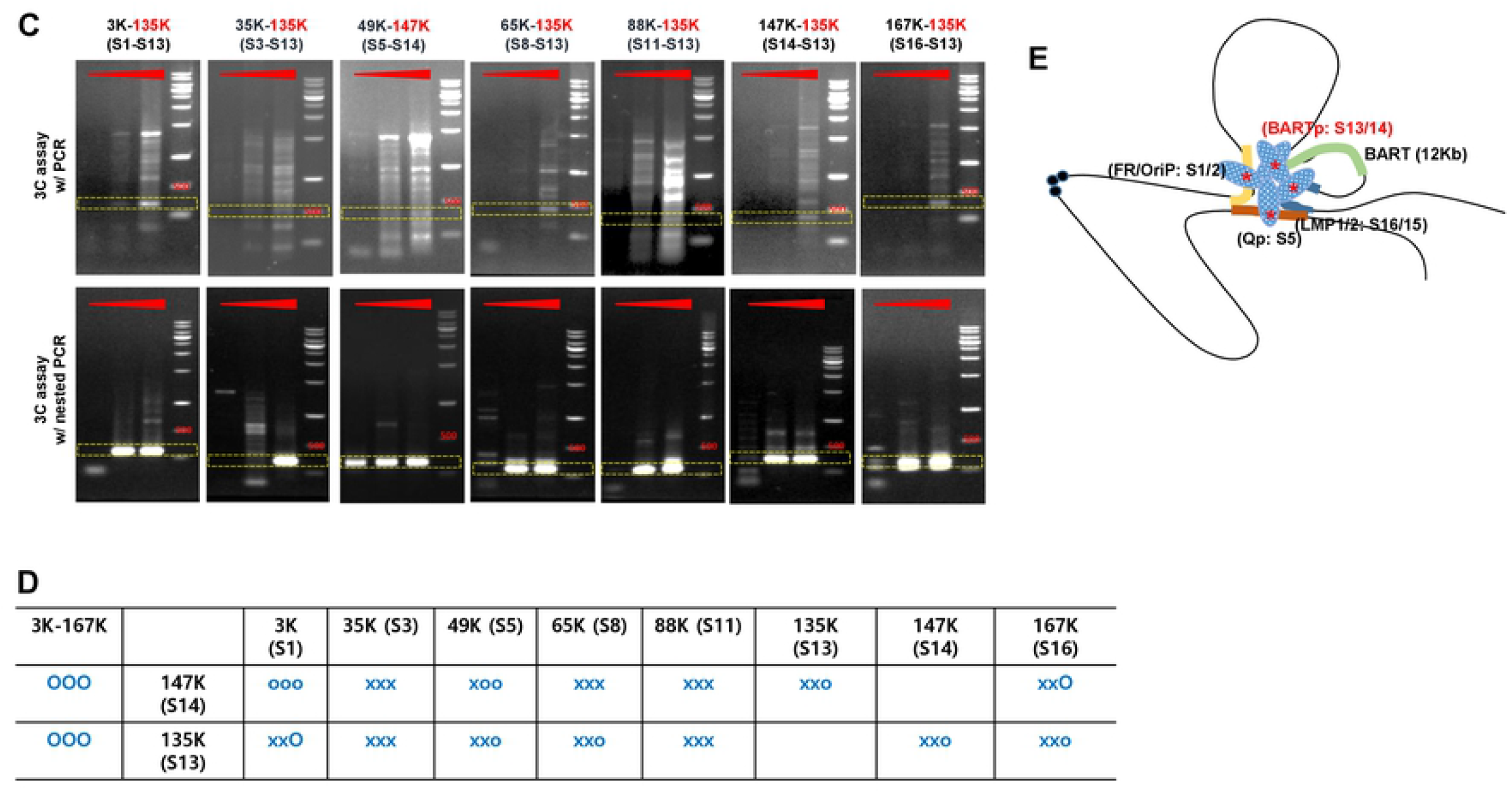
Confirmation of chromatin interaction by 3C-semi-quantitative (sq)PCR assay. **A)** Sites of *XhoI* restriction sites and 3C-sqPCR primers on EBV genome. **B)** 3C-sqPCR assay using 147K (S14) region as a view primer. EBV genome in SNU719 cells were cut by *XhoI* restriction enzyme, ligated by T4 DNA ligase, and then purified as described previously. Resultant EBV genome DNA was subjected to 3C-sqPCR assay. **C)** 3C-sqPCR assay using 135K (S13) region as a view primer. **D)** Summary of 3C-sqPCR assay data in table. O and o stand for strong and weak chromatin interactions, respectively. x stands for no detection. **E)** A simple diagram to indicate a DNA loop cluster mediated by BARTp CTCF BS associated chromatin interactions in SNU719 cells.

### Establishment of HEK293-EBV BART^−^ and BART^+^ BARTp CTCF BS smutant cells

EBV B(+) and B(-) (B(+/−)) genome sequences were compared in previous study [45]. This comparison revealed both loss of 11.8-kb BART region in EBV B(-) genome still retained the BARTp CTCF BS (S13) similar to EBV B(+) genome (Supplemental Fig. 1A). As BARTp CTCF BS is located in *RPMS1* intron region, an introduction of site-directed mutation in BARTp CTCF BS would not affect the *RPMS1* expression (Supplemental Fig. 1B). Resultant EBV B(+) S13^−^ bacmid from red-recombination was confirmed the site-directed mutation in BARTp CTCF BS by Sanger DNA sequencing (Supplemental Fig. 1C). In parallel, EBV B(-) S13^−^ bacmid was also confirmed the mutation by Sanger DNA sequencing (data not shown). Thereafter, EBV B(+/−) S13^+/−^ (B(+)S13^+^, B(+)S13^−^, B(-)S13^+^, B(-)S13^−^) bacmids were further tested their stabilities by EcoR1-digestion. Using these methods, we did not find any additional loss of EBV DNA (Supplemental Fig. 1D), nor defects in their ability to express GFP after several passages (Supplemental Fig. 1E).

### Effects of BARTp CTCF BS mutation on EBV infection

EBV B(+/−) S13^+/−^ (B(+)S13^+^, B(+)S13^−^, B(-)S13^+^, B(-)S13^−^) bacmid genomes were tested for their EBV infectivity to HEK293 cells. To this aim, HEK293-EBV B(+/−) S13^+/−^ cells were transfected with pcDNA3-BZLF1 and pcDNA3-BALF4. B(+/−) S13^+/−^ EBVs were harvested three days post transfection (Fig. 5A). Harvested viruses were tested their infectivity to HEK293 cells as mentioned above (Fig. 5B and 5C). Interestingly, both B(+) S13^−^ and B(-) S13^−^ EBVs were severely defected in their infectivity to HEK293 cells, compared to B(+) and B(-) S13^+^ EBVs. Furthermore, B(-) S13^+^ EBV persisted longer in HEK293 cells than B(+) S13^+^ EBVs. Taken together, these results indicated that BARTp CTCF BS, not the BART transcript, is required to maintain a full capacity of EBV infectivity.

**Fig 5.**
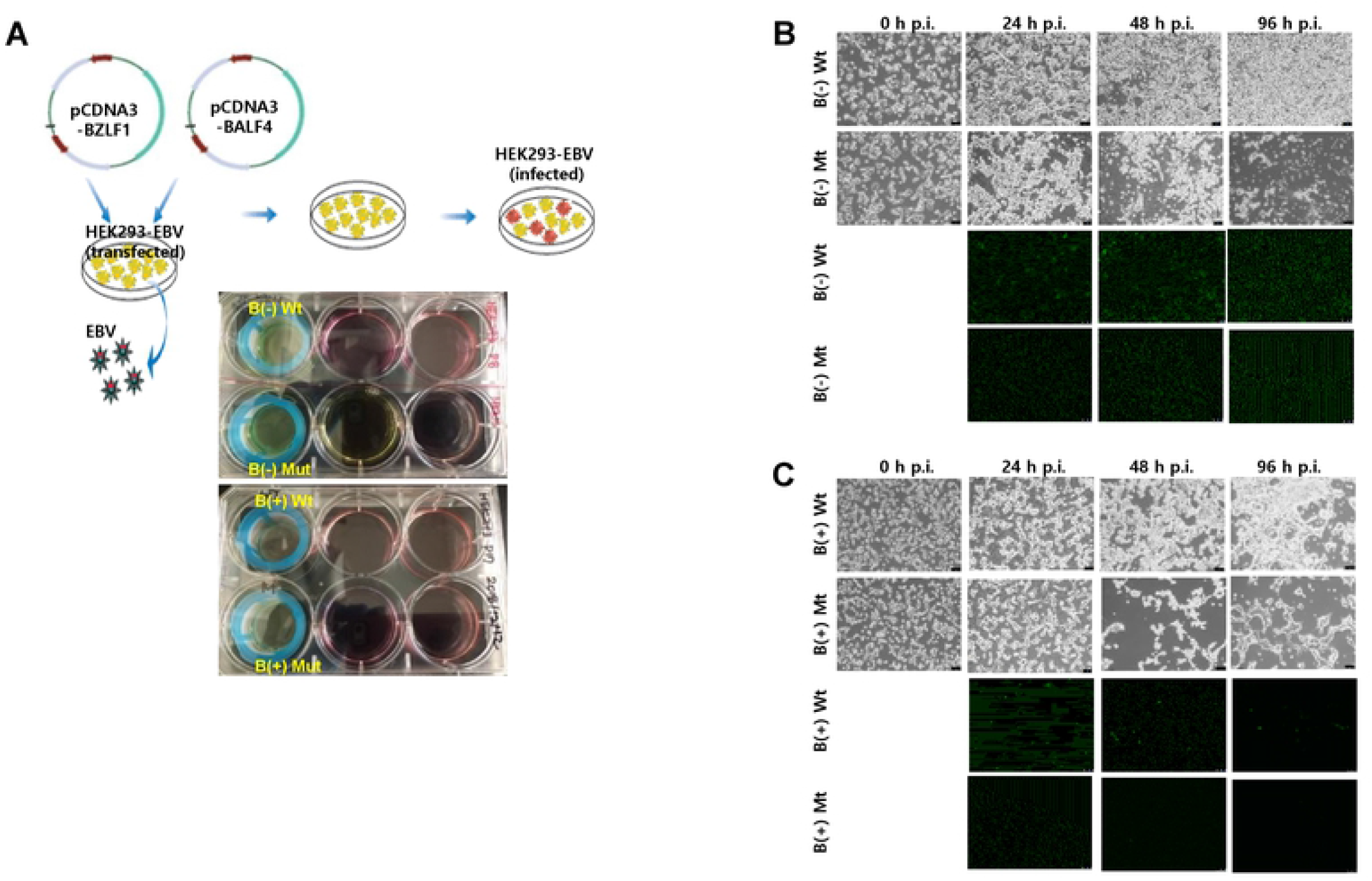
EBV infection assay. **A)** Schematic diagram of EBV infection assay. HEK293-EBV B(+/−) S13^+^ (Wt) and S13^−^ (Mt) cells were transfected with pCDNA3-BZLF1 and pcDNA3-BALF4. In three days post transfection, supernatants of transfected HEK293-EBV B(+/−) S13^+/−^ (Wt/Mt) cells were harvested and loaded on HEK293 cells freshly cultured on 6-well plate where cell strainer was equipped. EBV in harvested supernatants would infect HEK293 cells for 24 h. After infection, GFP from HEK293 cells infected by EBV was detected in series of time points. **B)** Infection assay of B(-) S13^+/−^ (Wt/Mt) EBVs to HEK293 cells. **C)** Infection assay of B(+/−) S13^+/−^ (Wt/Mt) EBVs to HEK293 cells.

### Effects of BARTp CTCF BS mutation on EBV gene expression

We further investigated overall expression patterns of EBV genes in HEK293-EBV B(+/−) S13^+/−^ (B(+)S13^+^, B(+)S13^−^, B(-)S13^+^, B(-)S13^−^) cells. The mRNA levels of *EBNA1, LMP2, BZLF1* were significantly lower in B(+/−) S13^−^ EBVs than B(+/−) S13^+^ EBVs (Fig. 6A and 6B). Consistently, the protein levels of *EBNA1, LMP2, BZLF1* were significantly lower in B(+/−) S13^−^ EBVs than B(+/−) S13^+^ EBVs (Fig. 6C and 6D). Intracellular *EBNA1* expression was also weaker in B(+/−) S13^−^ EBVs than B(+/−) S13^+^ EBVs which were visualized by immunofluorescence assay (Fig. 6E). Taken together, these results indicated that BARTp CTCF BS is required to maintain appropriate expression levels of EBV genes during their latent infection.

**Fig 6.**
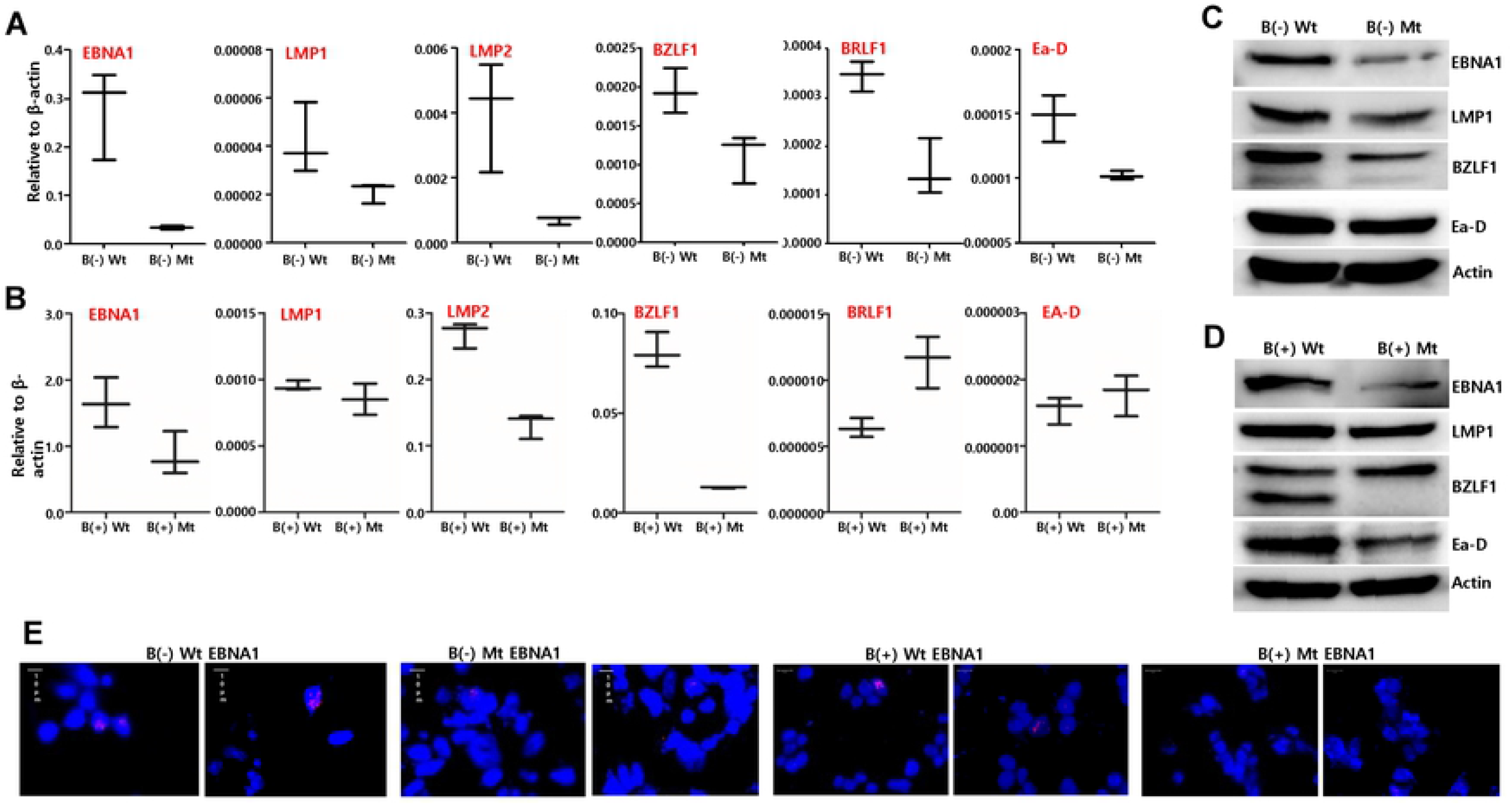
EBV gene expression in BARTp CTCF BS mutant EBVs. **A)** RT-quantitative (q)PCR assay was conducted to define profiles of EBV gene expression in HEK293-EBV B(-) S13^+/−^ (Wt/Mt) cells. **B)** The mRNA expression profile was defined in HEK293-EBV B(+) S13^+/−^ (Wt/Mt) cells. **C)** Western blot assay to compare EBV protein expressions between HEK293-EBV B(-) S13^+^ (Wt) and S13^−^ (Mt) cells. **D)** Western blot assay to compare EBV protein expressions between HEK293-EBV B(+) S13^+^ (Wt) and S13^−^ (Mt) cells. **E)** EBNA1 expressions in HEK293-EBV B(+/−) S13^+/−^ (Wt/Mt) cells were compared by immunofluorescence assay. Representive images for interphase nuclei stained with EBNA1 and Alexa flour 594 were shown.

### Effect of 11.8-kb BART region on CTCF enrichment around BARTp CTCF BS

Given the location of 11.8-kb BART region, we tested if 11.8-kb BART region can make regulatory effect to enrich CTCF on BARTp CTCF BS (S13) surrounding regions. For this aim, CTCF ChIP assay was performed to verify spatial effect of the BART region on the CTCF BSs to EBV genome. HEK293-EBV B(+/−) S13^+/−^ (B(+)S13^+^, B(+)S13^−^, B(-)S13^+^, B(-)S13^−^) cells were subjected to CTCF ChIP assay whose products were analyzed by real-time qPCR assays. As expected, CTCF was almost completely deprived from BARTp CTCF BS in B(+/−) S13^−^ EBVs (Fig. 7A and 7B). Interestingly, in other CTCF BSs surrounding BARTp CTCF BS, CTCF was differently distributed between B(+) S13^−^ EBV and B(-) S13^−^ EBV (Fig. 7A and 7B). Compared to B(+/−) S13^+^ EBVs, CTCF was further enriched at other CTCF BSs around OriP (S2) and LMP1/2 (S16) in B(-) S13^−^ EBV, while CTCF was severely deprived at those sites in B(+) S13^−^ EBV. These ChIP products were further analyzed to reconfirm CTCF distribution by semi-quantitative (sq) PCR assay (Fig. 7C and 7D). It was also observed that the enrichment and deprival of CTCF occurred at CTCF BSs around OriP and LMP1/2 dependent on absence and presence of BART, respectively. Taken together, these results from ChIP-(s)qPCR assays indicated that 11.8-kb BART region plays an important role in enriching CTCF at other CTCF BSs surrounding BARTp CTCF BS to regulate appropriate EBV genome 3D structure.

**Fig 7.**
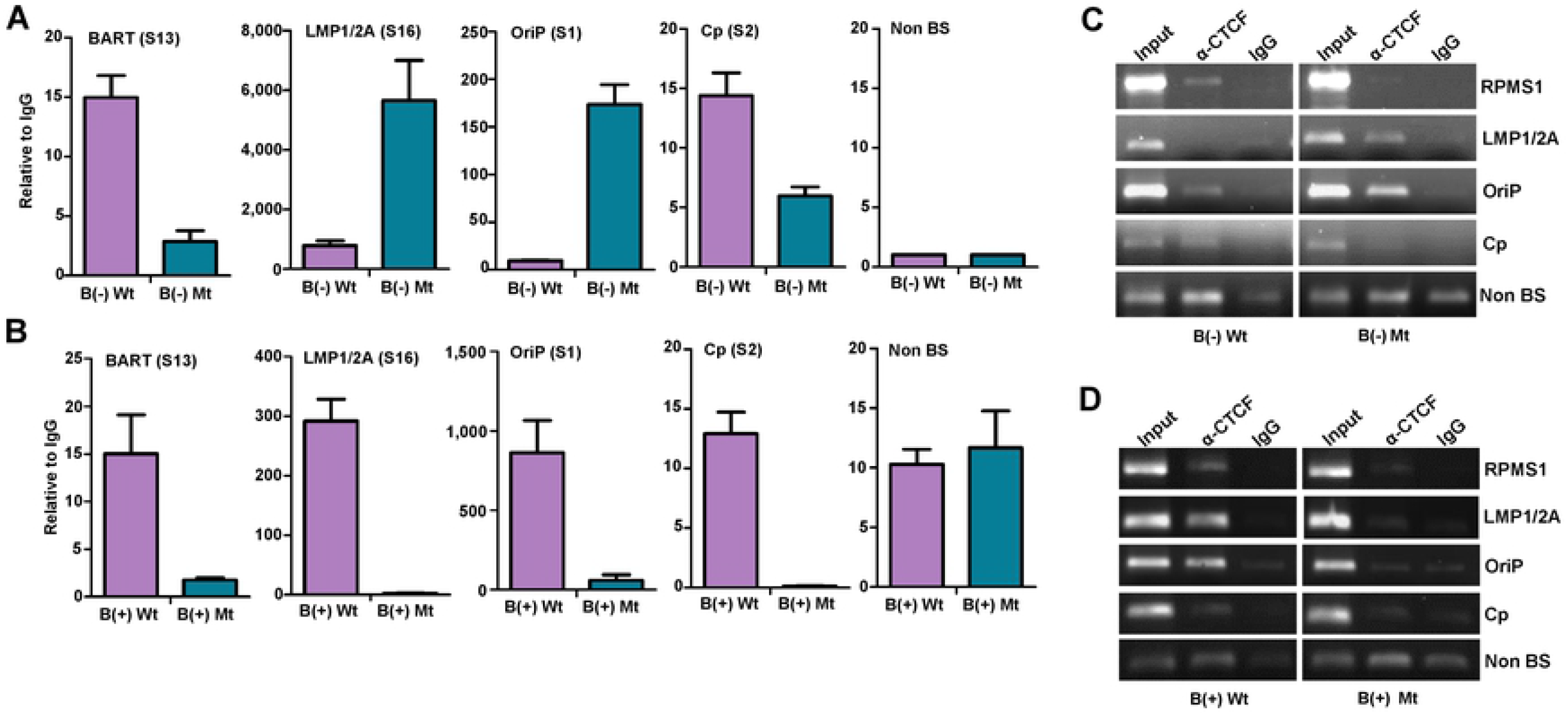
CTCF enrichment in HEK293-EBV B(+/−) S13^+/−^ cells. **A)** CTCF binding affinities were measured by CTCF ChIP-qPCR assay in HEK293-EBV B(-) S13^+/−^ (Wt/Mt) cells. **B)** CTCF binding affinities were measured by CTCF ChIP-qPCR assay in HEK293-EBV B(+) S13^+/−^ (Wt/Mt) cells. **C)** CTCF ChIP products from HEK293-EBV B(-) S13^+/−^ (Wt/Mt) cells were confirmed their enrichments using sqPCR. **D)** CTCF ChIP products from HEK293-EBV B(+) S13^+/−^ (Wt/Mt) cells were confirmed their enrichments using sqPCR assay.

### BARTp CTCF-mediates chromatin interactions

Since BARTp CTCF BS (S13) was implicated to differently distribute CTCF on its surrounding CTCF BSs, we tested whether BARTp CTCF BS might play a regulatory role in forming a cluster of several DNA loops mediated by BARTp CTCF BS. To this aim, 3C-sqPCR assays with HEK293-EBV B(+/−) S13^+/−^ (B(+)S13^+^, B(+)S13^−^, B(-)S13^+^, B(-)S13^−^) cells were conducted to consolidate chromatin interactions in similar way previously done with SNU719 cells in Fig. 4. Nuclei isolated from HEK293-EBV B(+/−) S13^+/−^ cells were subjected to 3C-sqPCR assay. Multiple chromatin interactions mediated by BARTp CTCF BS with loci such as OriP, Qp, and LMP1/2 were confirmed in HEK293-EBV B(+/−) S13^+^ cells (Fig. 8A). Like SNU719 cells, it was observed that 147K (S14) interacts with 3K (S1) and 167K (S16) regardless of 11.8-kb BART region. In spite of this similarity, both 167K-147K and 65K-147K interactions were slightly stronger in B(+) S13^+^ EBV and B(-) S13^+^ EBV. However, the site-directed mutation in BARTp CTCF BS caused to disrupt most chromatin interactions observed in B(+/−) S13^+^ EBVs (Fig. 8B). However, B(+/−) S13^−^ EBVs could not maintain almost the whole chromatin interactions directly associated with BARTp CTCF BS (Fig. 8B). The B(+/−) S13^−^ EBVs were not observed a specific chromatin interaction such as 65K (S8)-147K(S14). In spite of severe loss of chromatin interactions, some chromatin interactions indirectly associated with BARTp CTCF BS was relatively less affected in B(-) S13^−^ EBV. These indirectly associated chromatin interactions such as 3K-167K and 65K-147K were maintained in B(-) S13^−^ EBV although those interactions were abolished in B(+) S13^−^ EBV (Fig. 8B). All tested chromatin interactions were summarized in table (Fig. 8C) and depicted as simple diagrams (Fig. 8D). Given these data, BARTp CTCF BS (S13) could centralize key chromatin interactions among OriP, Qp, and LMP1/2. The 11.8-kb BART region could make structural effects on forming a key DNA loop cluster via BARTp CTCF BS-mediated chromatin interactions.

**Fig 8.**
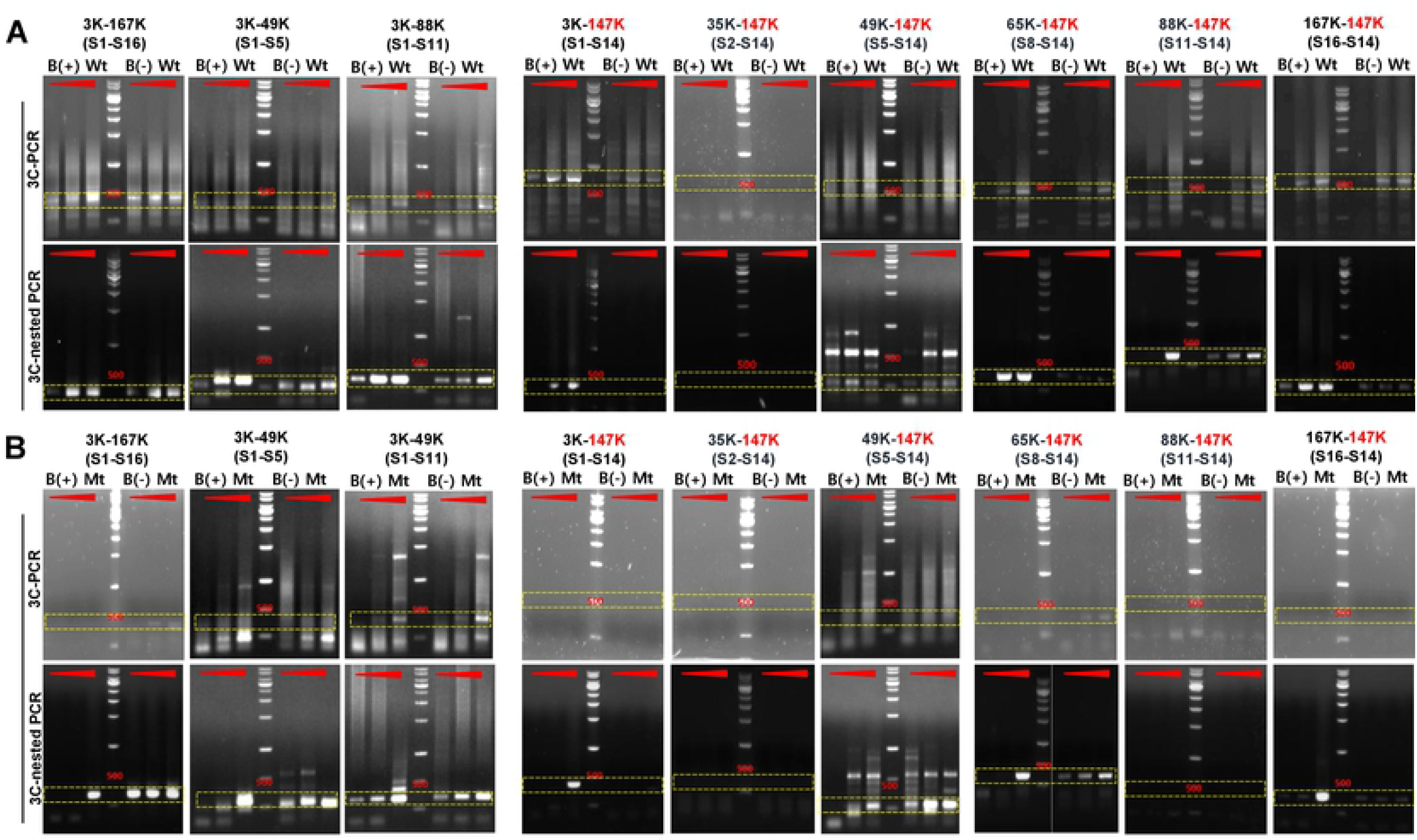
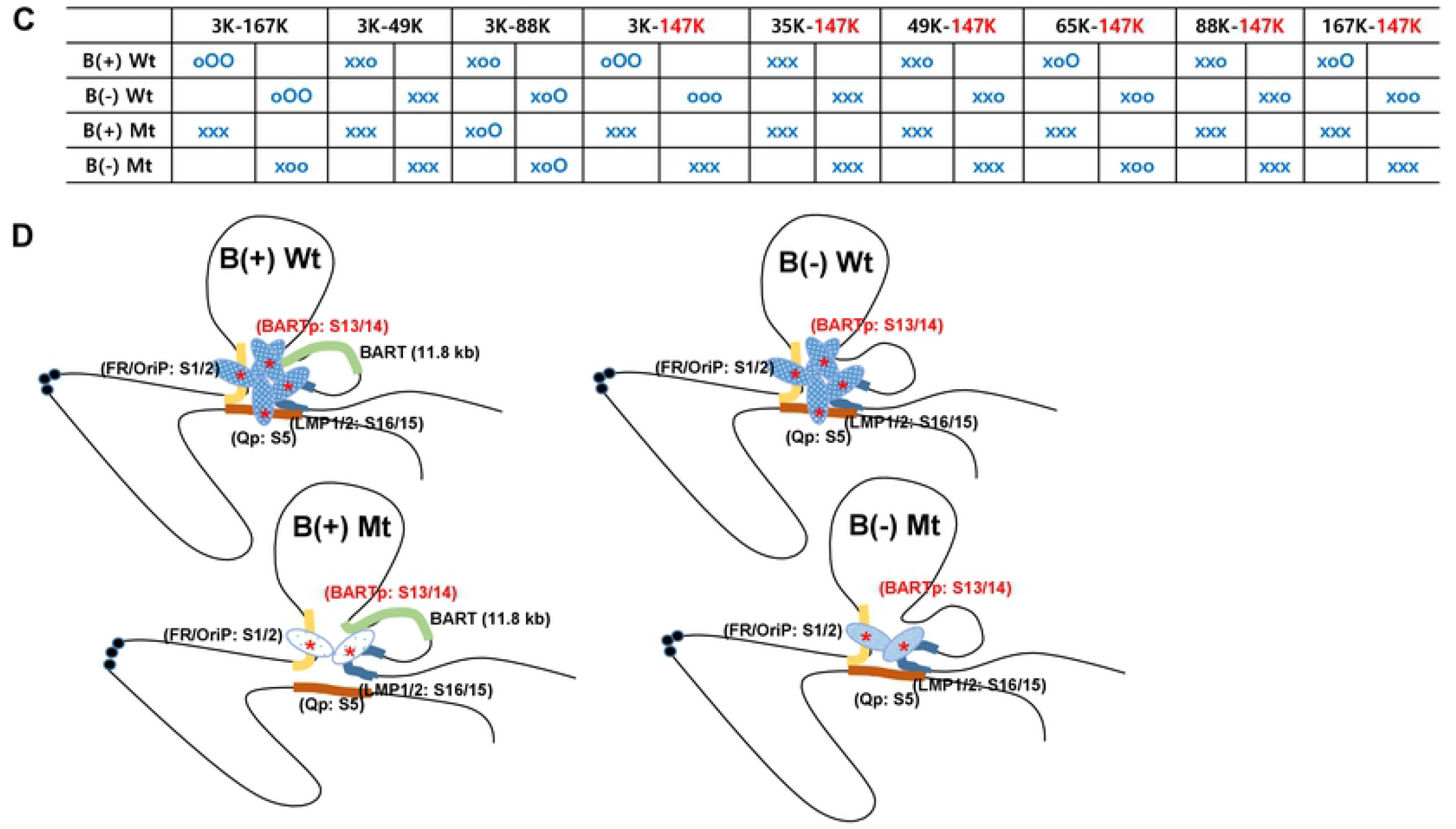
Confirmation of chromatin interaction affected by BARTp CTCF BS mutation. **A)** and **B)** Sites of *XhoI* restriction sites and 3C primers on EBV genome. EBV genome in HEK293-B(-) S13^+/−^ (Wt/Mt) EBV cells. were cut by *XhoI* restriction enzyme, ligated by T4 DNA ligase, and then purified as described previously. Resultant EBV genome DNA was subjected to 3C semi-quantitative PCR assay with view-point primers designed around 147K and 135K, respectively. **C)** Summary of 3C assays. O and o stand for strong and weak chromatin interactions, respectively. X stands for no detection. **D)** Simple diagrams to indicate a DNA loop cluster mediated by BARTp CTCF BS associated chromatin interactions in HEK293-EBV B(+/−) S13^+/−^ (Wt/Mt) cells.

## Discussion

Gene regulation requires integration of various signals and coordination of these signals across the genome. EBV gene expression is controlled at multiple levels, including transcription factor binding, transcription initiation and elongation, RNA processing, and epigenetic modification that control latent and lytic transcription [6]. In each of these context, CTCF-mediated chromatin interaction has played key roles in regulating EBV gene expression. CTCF BSs at *OriP* (S2 [46]), *Qp* (S5 [41]), and *LMP1/2* (S16 [43]) have been characterized their functional roles in regulating EBV gene expression. Most of these previous studies focused on EBV genomes in Burkitt’s lymphoma (BL) or lymphoblastoid cell lines (LCLs). Although 11.8-kb BART region was well conserved in EBV-associated Raji, Mutu, and GC1 cells, few genetic studies were conducted to define miRNAs in 11.8-kb BART region [38]. In particular, BARTp CTCF BS (S13) in gastric carcinoma cells has not been characterized its functional roles in regulating EBV gene expression.

Here, we first showed that BARTp CTCF BS (S13) can form a cluster of DNA loops via chromatin interactions mediated with several CTCF BSs such as *OriP, Qp*, and *LMP1/2*. Secondly, we found that functional BARTp CTCF BS (S13) is required to make a stable DNA loop between *OriP* and *LMP1/2*. Thirdly, we showed that BARTp CTCF BS (S13) is an important site of transcriptional regulation of EBV genes such as *EBNA1* and *BZLF1*. Finally, we found that functional BARTp CTCF BS (S13) is necessary for a full capacity of EBV infection. These findings suggested that BARTp CTCF BS (S13) coordinates EBV gene expression via forming interactions with other CTCF BSs in the viral genome.

CTCF has been previously implicated with cohesins in chromatin interactions by forming DNA loops [47]. Long-distance DNA interactions are essential to mediate communication between promoter and enhance elements. EBV OriP has been implicated as a transcriptional enhancer of *Cp* and *LMP1* promoter whose mechanism was to use the CTCF-mediated DNA loop structure [48]. In the present study, we examined the role of CTCF at the BARTp (S13) using EBV-associated gastric carcinoma cells and EBV infected HEK293 cells. We found that deletion of the BARTp CTCF BS (S13) in B(-) EBV genome resulted in more enrichment of CTCF at CTCF BSs in *OriP* and *LMP1/2*, while the deletion in B(+) EBV caused to deprive almost all the CTCF from CTCF BSs in *OriP* and *LMP1/2*. In similar context, the chromatin interactions of *OriP* and *LMP1/2* was more severely defected in EBV B(+) S13^−^ genome than EBV B(-) S13^−^ EBV genome due to the differential distribution of CTCF. These results suggested that 11.8-kb BART region could make spatial effects on forming CTCF-mediated DNA loops of *OriP* and *LMP1/2* loci.

Taken together, these results suggested that BARTp CTCF BS (S13) can play a complex role in regulating epigenetic modifications at both BART region and its surrounding regions such *LMP1/2* locus and *OriP* locus. One possible function of CTCF at the BARTp is to link *LMP1/2* locus, *OriP* locus, and *Qp* locus into a cluster of DNA loops. This genomic clustering would partly account for the role of CTCF in maintaining EBV latent infection. Similar to this present study, we previously observed that the chromatin interaction between *LANA* locus and *RTA* locus could occur to form a chromatin complex via CTCF during KSHV latent infection. In KSHV, these CTCF-mediated interactions were disrupted during KSHV lytic reactivation [49]. However, for EBV it is not yet known whether CTCF binding and genome conformation change during EBV latent-lytic switch.

CTCF has been implicated as a chromatin insulator and boundary factor [50, 51]. CTCF can prevent epigenetic drift by blocking heterochromatin formation at the EBV *Qp* region [41]. Deletion of CTCF BS in *LMP1/2* resulted in disrupting *OriP* and *LMP1/2* locus interaction, an increase in histone H3K9me3 and DNA methylation at LMP1 promoter region, and severe reduction of EBV latent infection [43]. In the present study, the deletion of CTCF BS in BARTp (S13) resulted in loss of *EBNA1* and *BZLF1* in mRNA and protein levels. In addition, the deletion caused to severely reduce EBV latent infection regardless of presence of 11.8-kb BART region. Thus, phenotype of BARTp CTCF BS (S13) mutant EBVs was similar to that of *LMP1/2* CTCF BS mutant EBV. One of possible mechanism is that BARTp CTCF might work together with *LMP1/2* CTCF to block the spread of heterochromatin that might be generated by GC-rich respective DNA of the EBV terminal repeat (TR) region. However, it remains to further study molecular mechanism that BARTp CTCF uses to coordinate with *LMP1/2* CTCF in blocking the spread of heterochromatin for EBV gene expression.

Recent studies using Hi-C methods examined EBV interactions with host chromosome in various cell types [40, 52]. Another related study used capture Hi-C to analyze KSHV looping DNA interactions during latency and reactivation, and found that DNA loops organized around highly active RNA polymerase II promoters, especially that for the viral PAN non-coding RNA [53]. In the present study, we used Hi-C method to define DNA looping interactions within EBV genome in EBVaGC and focused on the regions controlling the BART transcripts. Similar to KSHV PAN promoter, we found the EBV BART promoter to be an organizing hub for the EBV genome. While we did not examine RNA pol II binding, our findings indicate that CTCF-mediated chromatin interaction is likely to account for most DNA loops within EBV genomes. It is also possible that EBV association with host chromosome may also contribute to some aspects of EBV chromosome conformation. Although our high-resolution 4C analysis did reveal extensive conformational structure of EBV genome during EBV latency in EBVaGC, further studies will be required to resolve some of these more complicated functions of CTCF and to better understand how EBV has exploited CTCF binding sites to confer coordinate gene regulation and genome propagation in latent infection.

## Author’s Contributions

Conception and design: H. Kang, H, Cho, S. Development of methodology: H. Kang, K. Kim Acquisition of data (provided animals, acquired and managed patients, provided facilities, etc.): H. Kang, K. Kim, S. Cho, T. Kim, S. Huh, L. Kim Analysis and interpretation of data (e.g., statistical analysis, biostatistics, computational analysis): H. Kang, H. Cho, H. Tanizawa, J. Shin, Kyoung Jae Won Writing, review, and/or revision of the manuscript: H. Kang, H. Cho, P. Lieberman Administrative, technical, or material support (i.e., reporting or organizing data, constructing databases): T. Kanda, P. Lieberman Study supervision: H. Kang, H. Cho

---

**Supplemental Fig 1.**
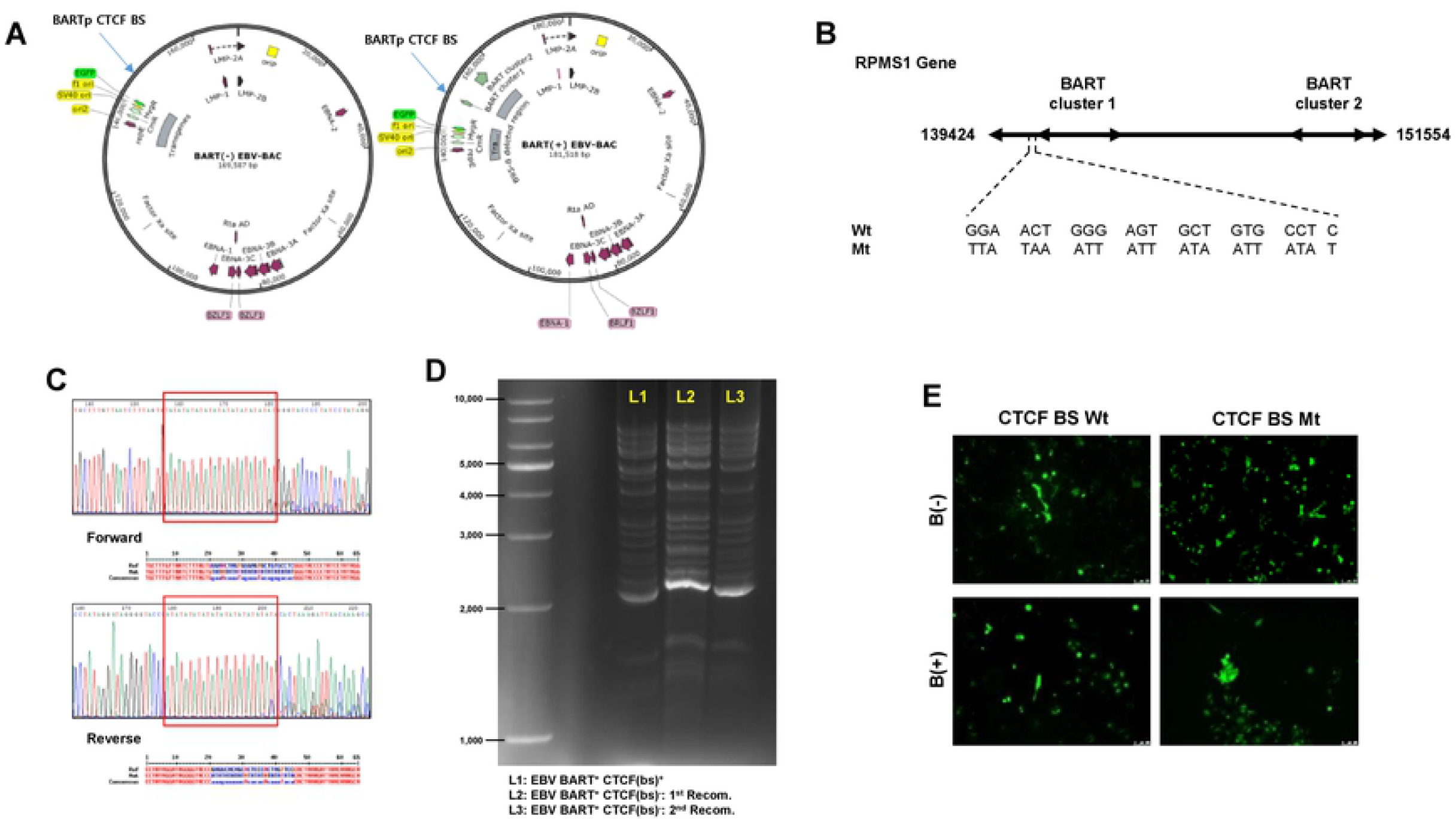
Construction of HEK293-EBV B(+/−) S13^−^ cells. **A)** Schematic diagram of EBV B(+/−) S13^+^ (Wt/Mt) bacmids. **B)** Sequences introduced at points of recombination to make site-directed mutations in BARTp CTCF BS. **C)** Confirmation of the site-directed mutation in BARTp CTCF BS in EBV B(+) S13^−^ (Mt) bacmid BY Sanger DNA sequencing. In parallel, EBV B(-) S13^−^ (Mt) bacmid was also confirmed the mutation by Sanger DNA sequencing (data not shown). **D)** Gel electrophoresis to check EBV genome stabilities of EBV B(+) S13^−^ (Mt) bacmids by *EcoRI* digestion; EBV B(+) S13^+^ (Wt) bacmid (lane 1), EBV B(+) S13^−^*bacmid-Kan^r^* with 1^st^ recombination (lane 2), and EBV B(+) S13^−^ (Mt) bacmid with 2^nd^ recombination (lane 3). We could not find any loss in EBV B(+) S13^−^ (Mt) bacmid. In parallel, EBV B(-) S13^−^ (Mt) bacmid was also confirmed their stabilities without any loss (data not shown). **E)** HEK293 cells were transfected with EBV B(+/−) S13^+/−^ (Wt/Mt) bacmids and selected using hygromycin B to establish HEK293-EBV B(+/−) S13^+/−^ (Wt/Mt) cells. GFP expression were determined in 40 days after hygromycin B selection and several passages. Established HEK293-EBV B(+/−) S13^+/−^ (Wt/Mt) cells were confirmed to maintain their GFP expressions even after several passages.

